# A domesticated fungal cultivar recycles its cytoplasmic contents as nutritional rewards for its leafcutter ant farmers

**DOI:** 10.1101/2022.03.03.482823

**Authors:** Caio Ambrosio Leal-Dutra, Lok Man Yuen, Pedro Elias Marques, Bruno Augusto Maciel Guedes, Marta Contreras-Serrano, Jonathan Zvi Shik

## Abstract

Leafcutter ants farm a fungal cultivar (*Leucoagaricus gongylophorus*) that converts inedible vegetation into food that sustains colonies with up to millions of workers. Analogous to edible fruits of crops domesticated by humans, *L. gongylophorus* has evolved specialized nutritional rewards—swollen hyphal cells called gongylidia that package metabolites ingested by ant farmers. Yet, little is known about how gongylidia form, and thus how fungal physiology and ant provisioning interact to farming performance. We explored the mechanisms governing gongylidium formation using microscopy imaging of ant-cultivated fungus and controlled *in vitro* experiments with the cultivar grown in isolation from ant farmers. First, *L. gongylophorus* is polykaryotic (up to 17 haploid nuclei/cell) and our results suggest intracellular nucleus distributions govern gongylidium morphology with their absence in expanding edges arresting apical growth and their presence mediating complex branching patterns. Second, nanoscale imaging (SEM, TEM) shows that the cultivar recycles its own cellular material (*e.g*. cytosol, mitochondria) through a process called ‘autophagy’ and stores the resulting metabolites in gongylidia. This autophagic pathway is further supported by gongylidium suppression when isolated fungal cultures are grown on media with autophagy inhibitors, and differential transcript expression (RNA-seq) analyses showing upregulation of multiple autophagy genes in gongylidia. We hypothesize that autophagic nutritional reward production is *the* ultimate cultivar service and reflects a higher-level organismality adaptation enabled by strict symmetric lifetime commitment between ant farmers and their fungal crop.

## Introduction

The advent of domesticated agriculture some 10 000 years ago was a turning point for humans and for the domesticated crops whose derived traits would likely have been maladaptive in their free-living ancestors [1–3]. Key crop adaptations include whole genome duplication events (resulting in polyploidy) that can increase functional heterozygosity [4] and selection for specific regulatory genes that can reduce seed shattering or enhance fruit size, color, and sweetness [5–8]. Fascinatingly, humans are not the only farmers. Several insect lineages independently evolved obligate farming systems of fungal cultivars that produce specialized edible reward structures [9]. However, while human farmers modify growth environments in well-known ways to maximize crop yield (e.g. adding fertilizers, controlling watering conditions, etc.), the analogous mechanisms by which insect farmers promote expression of edible reward structures in fungal cultivars remain poorly understood.

The largest-scale insect farmers are the *Atta* leafcutter ants, the crown group of the fungus-farming ‘attine’ ant lineage [10, 11]. Despite clear analogies with farming systems of humans [9], farming by leafcutter ants is fundamentally different because it is ‘organismal’ in the sense that it represents a strictly symmetrical obligate mutualistic dependence [12]. Such an arrangement usually does not allow alternative crops, but does sustain selection for co-evolutionary integration and higher-level adaptation that cannot evolve when farming practices are asymmetrically promiscuous [13, 14]. These differences make it interesting to explore how leafcutter farmers regulate crop productivity, since they can help to: 1) understand the broad eco-evolutionary success of these naturally selected farming systems and 2) provide nutritional insights into whether the leafcutter ectosymbiosis has achieved an organismal level of conflict-free trait evolution typically only seen in lifetime committed endosymbioses (*e.g.* the mitochondria-nucleus partnership in eukaryotic cells [15, 16]).

A mature rainforest colony of the leafcutter ant *Atta colombica* can have millions of specialized ants that divide the work of foraging fresh plant fragments and caring for the fungal cultivar [17]. In this way, colonies convert foraged fragments from hundreds of plant species [18] into a mulch used to provision their fungal cultivar *Leucoagaricus gongylophorus.* In return, the cultivar converts inedible plant biomass into edible reward structures called gongylidia, that are swollen hyphal cells *ca.* 30 μm in diameter and grow in bundles called staphylae [19–27].

Gongylidia are a defining trait of irreversible crop domestication and are unique to the fungal lineage farmed by leafcutter ants and other higher-neoattine genera including *Trachymyrmex, Sericomyrmex, Mycetomoellerius,* and *Paratrachymyrmex* [20, 28, 29].

Gongylidia mediate functional integration with their ant symbionts in two main ways. First, they contain enzymes (*e.g.,* laccases, pectinases, proteases) that ants ingest and then vector to patches of newly deposited vegetation to catalyze fungus-mediated digestion and detoxification [30–34]. Second, they contain nutrients (*e.g.*, amino acids, lipids and glycogen) that are the ants’ primary food source [35, 36]. The ability to regulate the quantity and quality of gongylidia would thus provide clear benefits for the ant farmers. However, the mechanisms linking substrate provisioning by farming ants and the production of the cultivar’s edible yield have remained poorly known. To better understand these mechanisms, we: 1) visualized the morphology of gongylidia and staphylae using scanning electron microscopy (SEM), 2) described the cellular reorganizations that mediate gongylidium formation by combining light, fluorescence, confocal and transmission electron microscopy (TEM), and 3) used a transcriptomics experiment to compare gene expression in hyphae and gongylidia to help resolve the metabolic pathways underlying gongylidia production.

We next examined the cellular origins of the edible resources contained in the large vacuole that fills each gongylidium cell. Previous evidence suggests that *L. gongylophorus* directly metabolizes provisioned plant fragments to produce these edible resources. First, the cultivar can metabolize lipids rich in alpha-linoleic acid (18:3) from foraged plant fragments into linolenic acid (18:2) that is enriched in gongylidia [37]. This synthesized metabolite is thought to mediate interkingdom communication by eliciting attractive behaviors in ant workers, in contrast to the precursor 18:3 lipid that elicits antagonistic behaviors [37]. Second, isotopic enrichment studies have shown that the cultivar quickly (within two days) shunts C and N from provisioned substrates (glucose and ammonium nitrate, respectively) into edible gongylidia [38]. Third, different substrate types are associated with increased expression of genes regulating targeted pathways for nutritional metabolism [21, 36, 39, 40]. However, it is also reasonable to predict that the diversity of compounds found within gongylidia have a diversity of biochemical origins.

Autophagy is a plausible alternative and/or complementary pathway underlying gongylidia formation that involves the recycling of the cultivar’s own metabolic source material and potentially the fine-tuning of its composition. The metabolic pathways for autophagy are conserved across eukaryotes and are known to mediate development, cellular differentiation [41–44], and housekeeping [45, 46] in fungal cells. During autophagy, the cultivar’s own cytoplasmic components (*i.e.*, glycogen, proteins, organelles) are incorporated into a vacuole for enzymatic degradation and the resulting catabolites are then recycled as nutrients to sustain other cellular processes and produce new cellular components [47, 48]. Initial evidence for autophagy in *L. gongylophorus* was first obtained in 1979 by Angeli-Papa and Eymé [49] who used TEM imaging to observe endoplasmic reticulum membranes engulfing mitochondria during gongylidium formation. However, to our knowledge, this preliminary evidence for autophagic recycling of the cultivar’s own intracellular content during gongylidium formation has not been subsequently explored.

We propose that confirmation of an autophagic pathway(s) would have important implications for understanding the leafcutter symbiosis since it implies that natural selection has targeted the farming symbiosis in ways that made provisioning more robust and less dependent on the variable quality and quantity of foraged vegetation. Specifically, we predict that autophagic nutrient recycling of cellular contents would: 1) reduce variability in the quality of the cultivar’s nutritional rewards and provide opportunities to optimize the composition of metabolic source material, 2) constrain the ability of ants to directly regulate cultivar productivity through their provisioning decisions, and 3) function in a complementary manner to provisioned plant substrates by providing a stable supply of metabolic precursor substrates during periods of environmental vegetation shortage.

We tested for autophagic gongylidium formation in three ways. First, autophagy encompasses two main types of cellular recycling mechanisms: 1) macroautophagy in which cytoplasmic content (i.e. cytosolic metabolites and organelles) are sequestered into double-membraned vesicles that fuse with vacuoles, and 2) microautophagy in which the vacuolar membrane invaginates and directly engulfs cytoplasmic cargo [47, 48]. We determined whether and how these autophagic processes influence gongylidium formation by examining organelle rearrangements in TEM images and tracking experimentally supplied fluorescent-labeled nutrients using confocal microscopy. Second, we tested whether autophagy is necessary for gongylidium formation by performing an *in vitro* experiment where the density of staphyla was measured in cultivars grown with known inhibitors and promoters of autophagy in fungal cells. Third, we tested whether autophagic pathways are differentially expressed in developing gongylidium cells by performing transcriptomic analyses of the mycelia and differentiated staphylae of the cultivar when grown under controlled *in vitro* conditions on a standardized medium without ant farmers.

## Results

### Nutritional reward structures

Each gongylidium cell consists of two sections that we term the bulb (swollen section) and the filament (elongated section) (Fig. 1 A). Gongylidia are often connected by intercalary bulbs (between filaments) and intercalary filaments (between bulbs) (Fig. 1 B-C) with multifaceted branching patterns and individual gongylidium cells bearing two or more terminal bulbs (Fig. 1 D). Gongylidia bulbs have variable diameters ranging from 12 μm to 50 μm and variable filament lengths ranging from 40 μm to > 250 μm. We hypothesize that variable bulb sizes reflect indeterminate growth trajectories of expanding gongylidium cells. All gongylidium cells also contained at least eight nuclei usually concentrated at the intersection of the bulb and the filament (Fig. 1 E) and usually one single vacuole (Fig. 1 F) that comprised up to half of each bulb’s total volume.

**Fig. 1:**
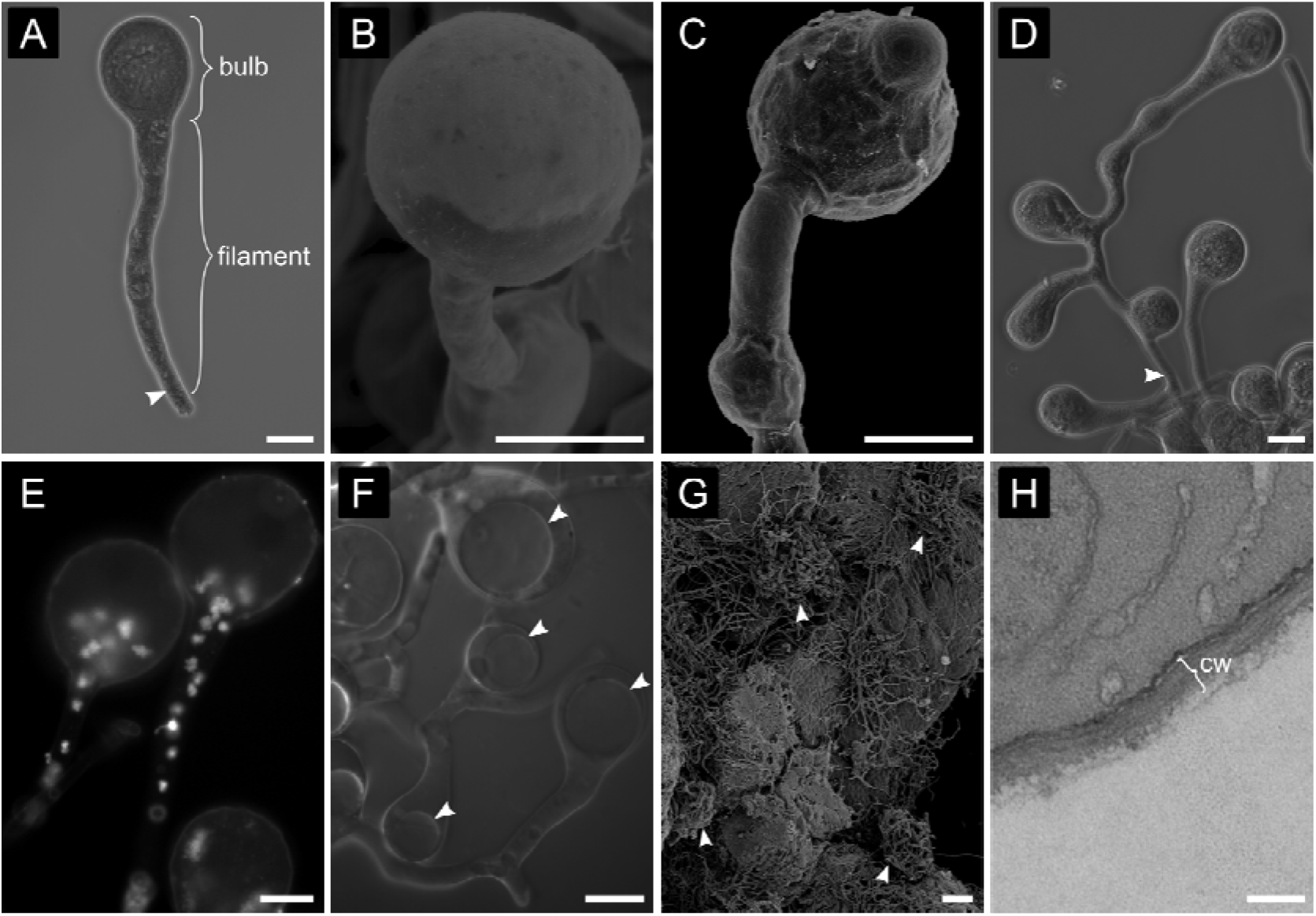
The *Leucoagaricus gongylophorus* fungal cultivar produces gongylidia as specialized nutritional reward structures for leafcutter ants. **A-C)** Gongylidium cells are typically depicted as a bulb at the end of a filament in the apical hyphal compartment separated by a septum (arrowheads). **D)** Gongylidia frequently exhibit more complex branching, with bulbs between filaments or in lateral branches of single hyphal cells delimited by septa. **E)** Individual gongylidium cells are polykaryotic [86], meaning that they have many haploid nuclei (white dots). Here, we show that in mature non-branching gongylidium cells, these nuclei occur at the base of the bulb (below a single large vacuole) and in the filament. Nuclei were visualized using DAPI staining. **F)** Each gongylidium cell contains a large vacuole (arrowheads). **G)** Staphyla grow in discrete patches at the surface of the fungus garden matrix in the middle garden stratum (arrowheads). **H)** Gongylidium cells have thin cell walls ranging from 120 to 220 nm (cw). Images produced by light microscopy (panels A, D, F), fluorescence microscopy stained with DAPI (panel E), SEM (panels B, C, G) and TEM (panel H). Scale bars: A-F = 20 μm, G = 100 μm, H = 200 nm.

Individual staphylae range widely in size, with tens to hundreds of individual gongylidium cells (Fig. 1 G, Fig. S1), and always formed on the surface of fungus garden mycelial matrix where ants can easily detach them from surrounding hyphae. Staphylae also form in the absence of ants under *in vitro* (Petri dish) growth conditions, but they have the following key morphological differences compared to those growing in ant-tended fungus gardens, being: 1) less detachable because they are usually covered by filamentous hyphae, 2) larger in area and with more individual gongylidium cells, and 3) comprised of several ‘burst’ gongylidium cells. We thus propose that under farming conditions, ants harvest staphylae earlier in their development before vacuoles can produce turgor pressure exceeding the retaining capacity of their exceptionally thin (ca. 120 to 220 nm) cell walls (Fig. 1 H).

### An autophagic mechanism of gongylidia formation

Microscopy images (TEM, light, fluorescence and confocal) of gongylidium cells revealed structures that are diagnostic of macroautophagic processes. First, gongylidia were enriched with long stretches of endoplasmic reticulum that produce double-membraned vesicles called autophagosomes (Fig. 2), within which recycling of cellular materials is initiated. We confirm that autophagosomes contained cytosol (Fig. 2 A), glycogen (Fig. 2 B) and mitochondria (Fig. 2 C), and predict that other metabolites (*e.g*., lipids, amino acids, enzymes [28, 30, 37, 40, 64]) are likely abundant, but are too small to be detected with TEM. Second, large numbers of damaged mitochondria were present in gongylidia and were often associated with endoplasmic reticulum membranes (Fig. 2 C) where they were likely destined to be sequestered into autophagosomes, digested, and recycled into edible metabolites. Third, vacuoles within gongylidium bulbs often contained single-membraned autophagic bodies (Fig. 2 D-F). These vesicles indicate the delivery of metabolites into vacuoles, since they are autophagosomes that lost their outer membrane after vacuolar fusion. This was confirmed by confocal images showing that autophagic bodies within gongylidium vacuoles contained fluorescently labeled sugars from the cultivar’s cytosol (Fig. 2 E-F, <u>Video S1</u>). Given these hallmarks of macroautophagy (and the lack of evidence for microautophagy), we henceforth use ‘autophagy’ to refer to macroautophagy.

**Fig. 2:**
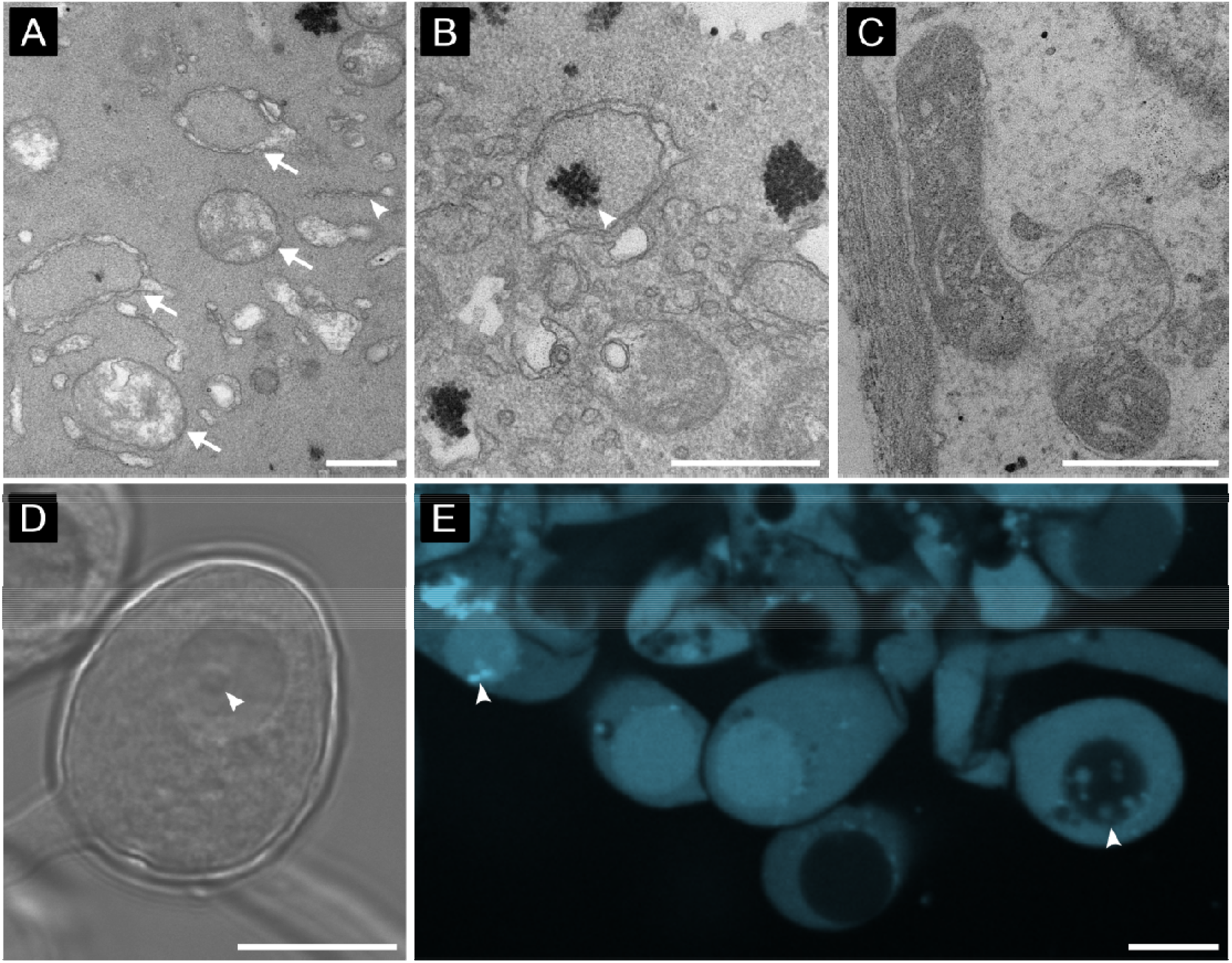
Autophagic recycling of cellular material. **A)** The autophagosomes that are diagnostic of autophagy are vesicles with double-layered membranes (arrows) and are produced by stretches of endoplasmic reticulum membranes (arrowheads). **B)** Autophagosomes sequester cytoplasmic components including glycogen (arrowhead) and **C)** mitochondria which are then delivered to vacuoles in gongylidia bulbs for further degradation. **D-E)** Vacuolar expansion is mediated by autophagosomes that lose their outer membrane after fusing with the vacuole. These single-membraned autophagic bodies are vesicles that can be seen inside the vacuole prior to their degradation (arrowhead). Images acquired by TEM (A-C), phase contrast microscopy (D) and confocal microscopy stained with dextran-Alexa fluor 647 (E). Scale bars A-C = 500 nm, D-E = 20 μm.

An autophagic mechanism for gongylidium formation was further supported by significant *in vitro* treatment effects of chemical autophagy inhibitors on staphyla density (Kruskal-Wallis: H_3_ = 34.7, *p* < 0.001, Fig. 3). Pairwise post-hoc comparisons indicated that both autophagy inhibitors (3-MA and CQ) had significantly reduced staphyla density compared to both the control (PDA) (*p_adj_* < 0.001 and *p_adj_* = 0.001, respectively) and the autophagy-promoter (RAP) treatment (*p_adj_* < 0.001 and *p_adj_* = 0.026, respectively). There was also a significant treatment effect on mycelial growth area (H_3_ = 16.3, p = 0.001). However, this was due to differences between 3-MA and all other treatments (PDA:3-MA, *p_adj_* = 0.004; RAP:3-MA, *p_adj_* = 0.015; CQ:3-MA, *p_adj_* = 0.001), with no other significant pairwise comparisons (Fig. S2). Thus, while both autophagy inhibition treatments resulted in staphyla reduction, it is possible that 3-MA negatively influenced staphyla density through unknown indirect effects on cultivar performance. The autophagy promotor (RAP) did not significantly increase staphyla density relative to control (PDA) or either of the autophagy inhibition treatments (Fig. 3).

**Fig. 3:**
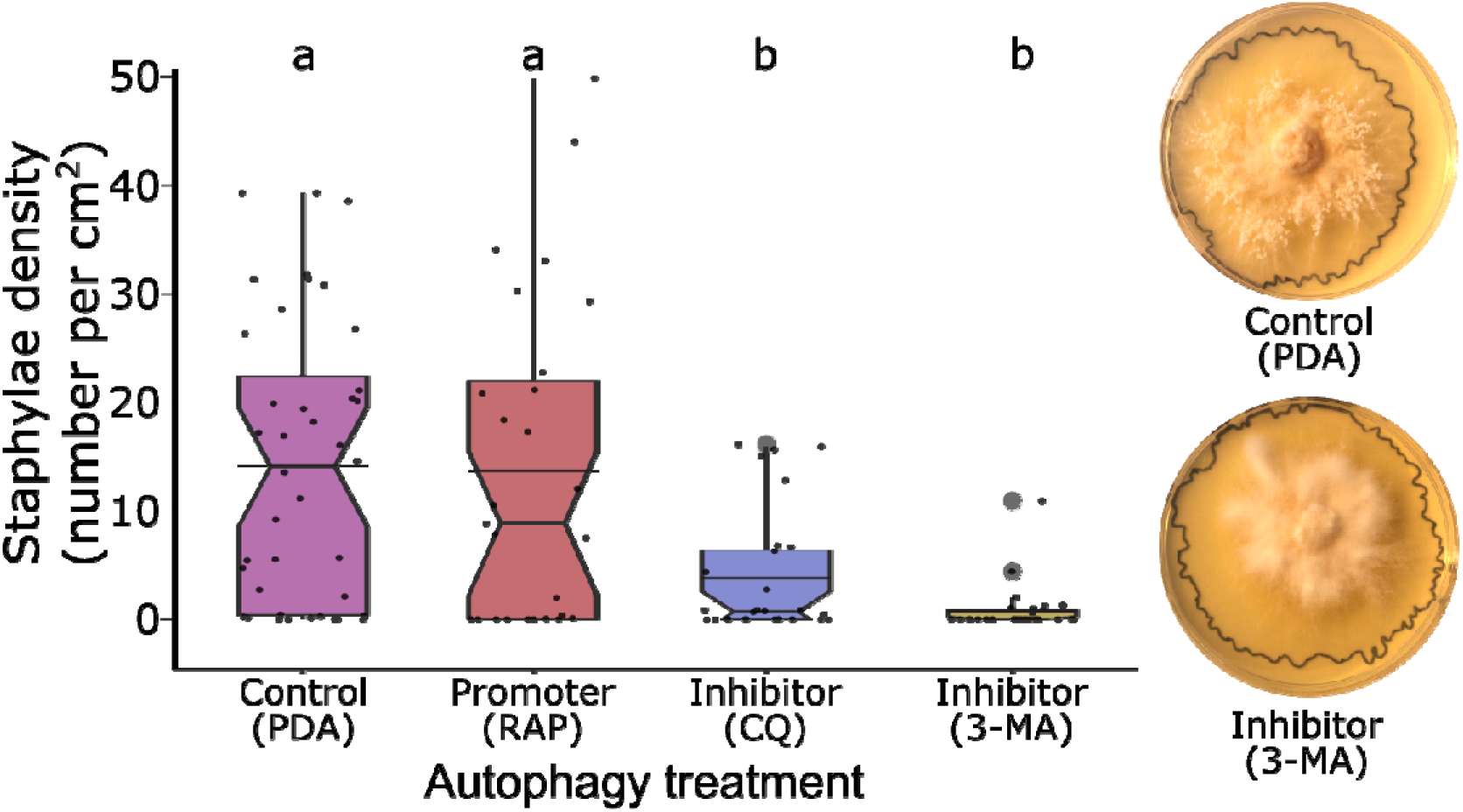
Experimental evidence that autophagic recycling of the fungal cultivar’s own cellular material mediates gongylidia formation. Gongylidium density was significantly inhibited when *L. gongylophorus* was grown on potato dextrose agar supplemented with one autophagy inhibitor chloroquine (CQ, n = 28) or 3-methyladenine (3-MA, n = 30)) relative to control (PDA, n = 40) and an autophagy promoter rapamycin (RAP, n = 27)). Representative Petri-dishes with control (PDA; top) and inhibition (CQ; bottom) are displayed at the right, black outlines indicate the radial growth area of cultivars and white fungal clusters in the control are the staphylae. Different letters above the boxes indicate significant differences determined by a post-hoc Dunn’s pairwise test (*p* < 0.05) and horizontal bars indicate the distribution means.

### Transcriptome assembly and autophagy pathway analysis

Using a *de-novo* assembly and a clustering analysis by similarity, we recovered 78,820 transcripts from the *L. gongylophorus* transcriptome of which, 4,755 were differentially expressed (log2 fold-change > 1.0, Benjamini–Hochberg adjusted *P* < 0.001) in staphylae (n = 3,011 transcripts) or in non-differentiated mycelia (n = 1,744 transcripts). Of these differentially expressed transcripts (DETs), 31 were assigned a KEGG orthology term associated with the yeast autophagy pathway (n = 22 in staphylae, n = 9 in mycelia, Table 1). Several key genes were upregulated in staphylae that are typically over-transcribed during autophagy [48]. These include genes (ATG7, ATG8, ATG10) linked to the recruitment of cargo into incipient unclosed membranes (*i.e.*, phagophores) and mature autophagosomes, as well as genes related to starvation signaling (Sch9, Tap42, ATG13 and RIM15), vacuole fusion machinery (Ypt7, Mon1 and Vps3), and degradation of autophagic bodies (PRB1 and ATG15) (Table 1, Fig. S3). In contrast, the few autophagy-specific genes upregulated in undifferentiated mycelia appear related to the starvation signaling step of autophagy induction, rather than a fully functioning autophagy pathway.

**Table 1:**
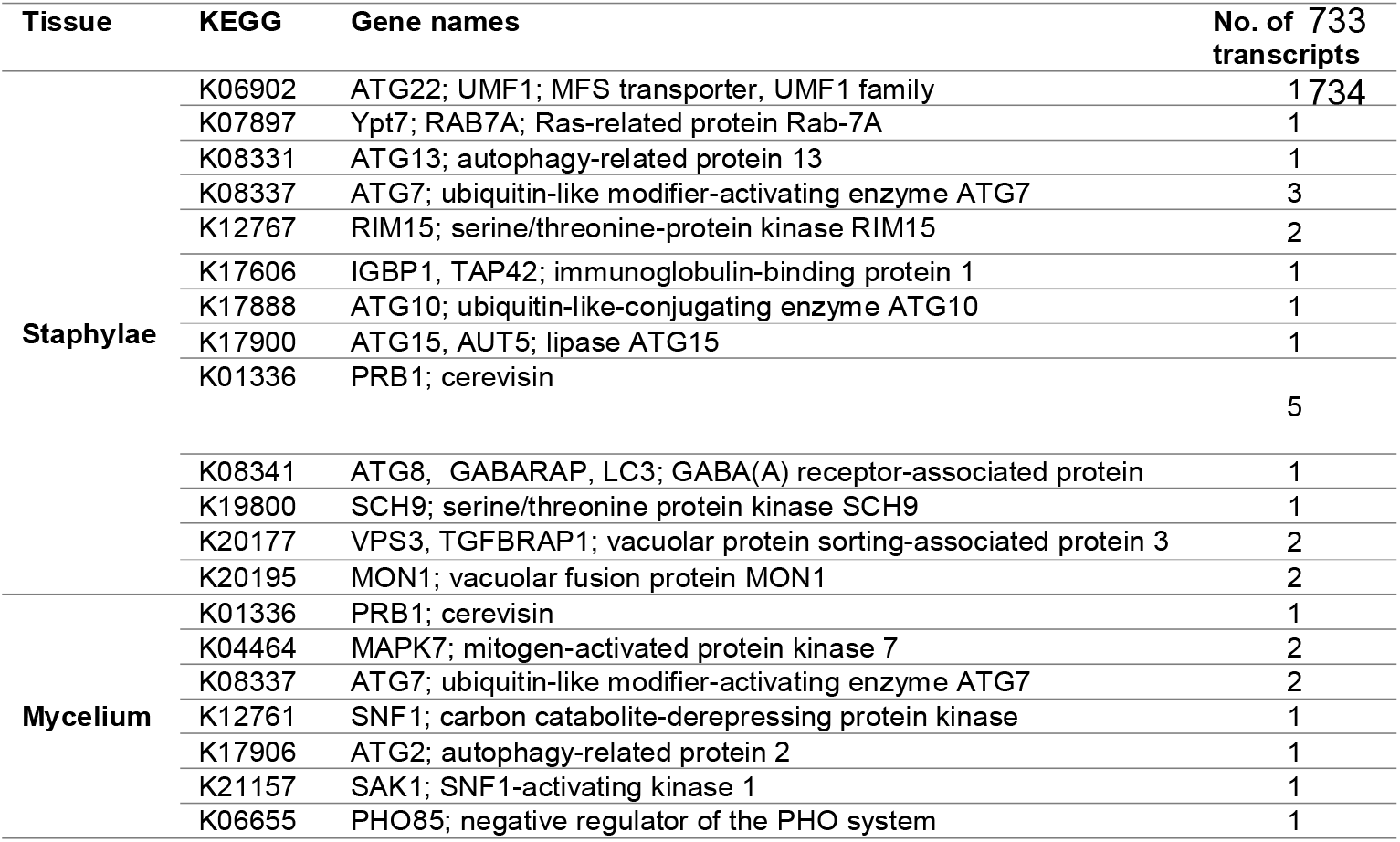
Upregulated transcripts in autophagy pathway annotated with KEGG database (see Fig. S3).

## Discussion

Our results provide novel insights into the mechanisms of higher-level homeostatic integration in a uniquely ‘organismal’ insect-microbe ectosymbiosis. Several lines of evidence indicate that the fungal cultivar of leafcutter ants uses autophagic recycling to convert its own cellular material into edible metabolites within specialized nutritional reward structures. First, nanoscale imaging shows the cellular hallmarks of autophagy (*e.g.* autophagosomes, autophagic bodies, abundant endoplasmic reticula) and indicates rapid delivery of labeled cytosol nutrients into gongylidium vacuoles (ca. 30 minutes). Second, experimental suppression of autophagy suppresses gongylidium density. Third, we find an upregulated autophagy pathway associated with differentiated gongylidium cells. We hypothesize that this autophagic recycling pathway represents a final domestication step where the cultivar came to unambiguously prioritize nutritional services to its hosts even at the expense (up to a point) of its own mycelial health. In this sense, the autophagic recycling pathway expresses an obligately symmetric commitment between symbionts achieved after the domesticated fungal cultivar lost the capacity for a free-living existence and became fully integrated into the host colony’s germ line.

We further propose that autophagic recycling facilitates homeostasis at higher levels of organization by stabilizing the quantity and nutritional quality of the cultivar’s nutritional rewards in fluctuating environments. First, the seasonal and spatial availability of preferred plant fragments may vary in suboptimal ways [65, 66]. Second, foraged plant fragments can contain key nutrients—but these nutrients can occur in suboptimal ratios and concentrations relative to the cultivar’s intrinsic needs and tolerances [19]. Third, plant fragments contain a wealth of recalcitrant compounds (e.g. cellulose and lignin) and toxic metabolites that can reduce cultivar performance [67–70]. During such periods of plant-fragment shortage [71–73], the cultivar may use autophagic recycling of its own organelles to yield reliably available and chemically predictable metabolic precursor compounds. Analogous adaptations aimed solely regulating homeostasis at higher levels of organization are absent from all other ectosymbioses that are promiscuous by comparison, where the interests of symbionts are not completely aligned, and partners must screen, sanction, and police to dissuade cheating [74–76].

The discovery of autophagic recycling also provides a new lens to interpret well-known gardening behaviors in leafcutter ants. For instance, gardening ants constantly prune the cultivar’s fungal mycelia and this has been hypothesized to cause mechanical disruptions that stimulate gongylidium formation [77]. We propose that such pruning behaviors sever hyphal connections and block the flow of nutrients to newly isolated fungal cells, which in turn induces gongylidium formation by causing an autophagic response to starvation—which is a common driver of autophagy in cells [50]. Additionally, previous *in vitro* studies have observed highest staphylae densities at the lowest nutrient concentrations suggesting a link between staphyla formation and nutrient depletion [19, 40, 78–80]. Thus, while much research assumes that cultivar production hinges on a maximized flow of provisioned plant fragments [19, 53, 81], the behaviors linked to targeted nutritional suppression may also be important for the production of nutritional rewards.

While gongylidia-linked autophagic recycling appears common, it likely complements other nutritional reward production mechanisms. First, leafcutter ants frequently ingest gongylidia contents (crushing the gongylidia to ingest their cellular content) and vector the cultivar’s enzymes [31, 32, 34, 35] and nutrients [35, 36] as fecal droplets to catalyze degradation and detoxification of newly deposited plant fragments [30]. Metabolites within these fecal droplets are assimilated by the cultivar and some subset likely enters biosynthetic pathways linked to gongylidium formation. Second, key nutrients also appear to derive from bacterial symbionts (rather than plant fragments or autophagic recycling) [82–84]. For instance, attine ants have lost the ability to synthesize arginine [80]—and depend on the cultivar’s metabolism to produce this nitrogen-rich amino acid [28]. In turn, the ants have evolved tight mutualistic associations with specialized bacterial symbionts that convert excess arginine in the ants’ guts into ammonia (the Mollicutes EntAcro1, [84]) that is an N-rich fertilizer vectored by the ants back to their fungal symbiont. As further evidence that the autophagic-recycling is one of several mechanisms by which gongylidia fill with metabolites, we observed that staphyla production was still possible (even though significantly reduced) when autophagy was inhibited in the *in vitro* experiment. Resolving whether and how these nutritional pathways fluctuate relative to the specific resource needs of the colony thus represents an important next step.

### Reconstructing the cellular reorganizations enabling gongylidium formation

The combined evidence we present throughout this study enables us to propose a complete pathway for gongylidia development (Fig. 4). Tissue differentiation initiates when a hyphal cell growing at surface interstices within the fungus garden matrix widens (Fig. 4 A) and continues as a vacuole begins expanding in response to autophagic recycling. This process accelerates autophagosomes fuse with the vacuole and delivers cytoplasmic cargo (Fig. 4 B). The location of this vacuole shapes the morphology of developing gongylidia because it mediates the spatial distributions of the up to 17 [86] haploid nuclei within individual gongylidium cells by excluding them from the apical tip (Fig. 4 C).

**Fig. 4:**
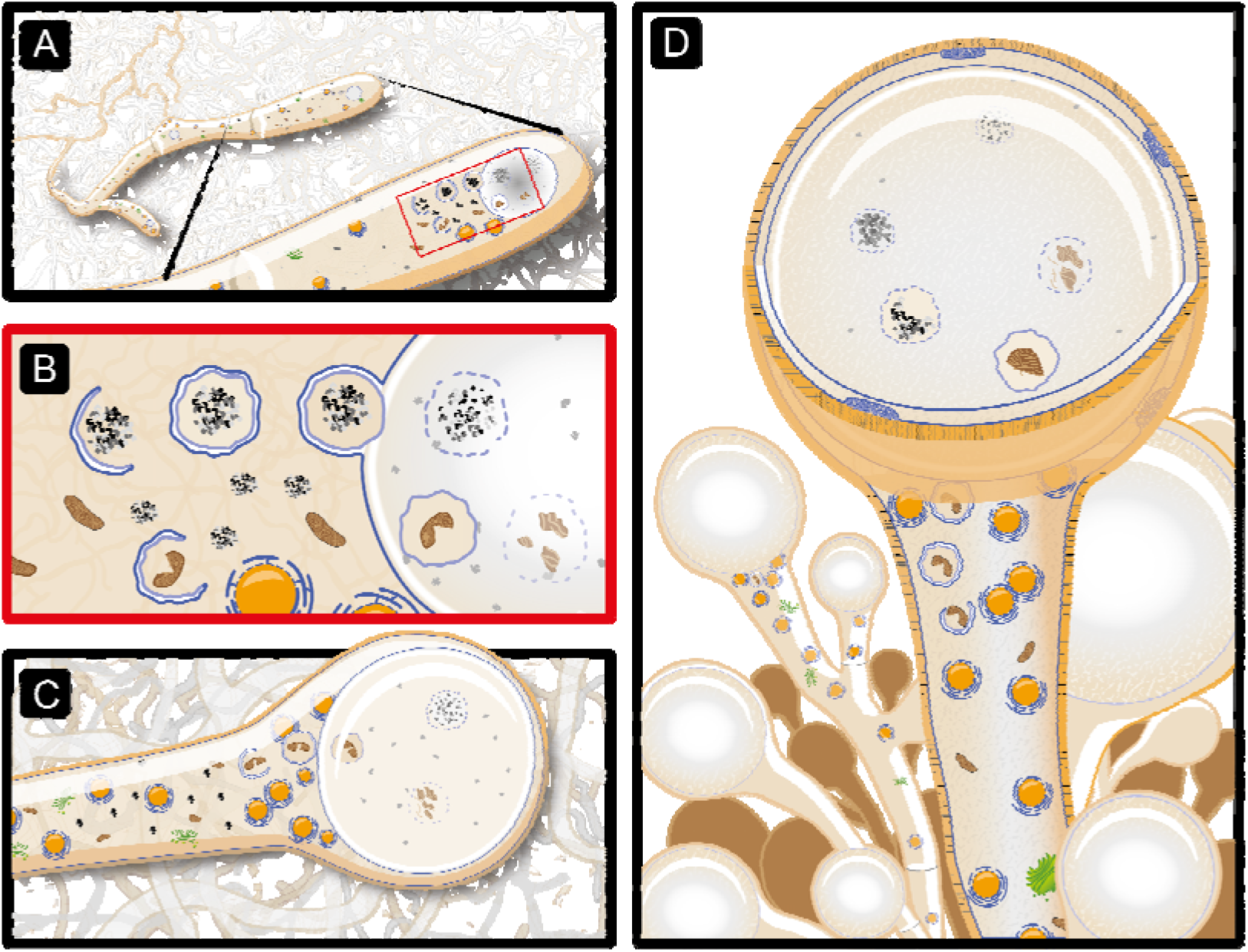
The hypothesized stages of autophagy-mediated gongylidium development. **A)** An unknown mechanism (potentially starvation mediated by ant pruning [77]) triggers the widening of ordinary hyphae. As the hypha elongates, the nuclei (orange circles) naturally migrate towards the hyphal tip. **B)** Mediated primarily by an autophagic process, a large vacuole expands with the fusion of newly formed double membrane vesicles called autophagosomes (blue membraned vesicles) that sequester material present in the cytosol like glycogen (black and gray aggregates) and damaged mitochondria (brown indented ovals). This process is indicated by the proliferation of endoplasmic reticulum membranes (blue membranes around nuclei) that produce autophagosomes. **C)** The fusion of autophagosomes into vacuoles mediates their expansion and either forces the apical bulb swelling while also halting further apical growth by excluding nuclei from the hyphal tip or produce an intercalary bulb when the vacuole is located among nuclei. **D)** This process repeats in up to hundreds of adjacent hyphae that become tangled to form the staphyla.

We hypothesize that the obstruction of nuclear migration causes the bulb’s balloon-like expansion by blocking communication between nuclei and the Spitzenkörper—the centralized machinery for hyphal growth located in the hyphae tip. At the transcript level, a structurally modified and upregulated transcript carrying a domain associated with microtubule related proteins [40] may mediate such nuclear migration in association with motor protein complexes [87–89]. This mechanism would resemble growth dynamics in most filamentous fungi where nuclei are distributed evenly throughout the hyphal compartment and promote tip elongation by migrating apically [88, 90]. Further supporting this hypothesis, when nuclei are occasionally aggregated in different regions of the filament and bulb, they are associated with ramified branching gongylidia and intercalary bulb formation. Finally, we propose that staphylae arise from this patchy ramification of tangled gongylidia (Fig. 4 D).

These results can also inform our understand of the functional consequences of polyploidy in the domesticated *L. gongylophorus* cultivar which contains up to 7 distinct haplotypes per cell [86]. Specifically, key next steps involve moving beyond distributions of nuclei in gongylidium cells, to testing whether factors like nucleus-specific expression and nuclear dominance are linked to gongylidium formation. Such a mechanism has recently been observed in the production of edible reward structures produced by the heterokaryon human-domesticated champignon fungus (*Agaricus bisporus*), where two distinct nuclear types exhibit differential expression in distinct tissues during mushroom formation [91]. The convergent existence of such molecular mechanisms in gongylidium formation would provide a further means of testing the hypothesis that these nutritional rewards are derived from cystidia [20], which are modified hypha found in the hymenium of several groups of basidiomycetes [92, 93]. Autophagic induction of cystidia may have provided crucial pre-adaptations harnessed by natural selection to generate the unique gongylidium reward structures.

## Methods

### Sample acquisition and fungal symbiont isolation

Two colonies of *Atta colombica* (Ac2012-1 and Ac2019-1) were used in this study that were collected in Soberanía National Park, Panama and thereafter maintained at the University of Copenhagen in a climate-controlled room (25°C, 70% RH, minimal daylight). For microcopy and imaging, we used staphylae collected directly from the colonies’ fungus gardens, and for the autophagy inhibition experiments and transcriptome sequencing, we used axenic cultures isolated from the fungal cultivar (*L. gongylophorus*) grown in 90 mm Petri dishes filled with 20 ml potato dextrose agar (PDA) that were kept in the dark at 25°C.

### Imaging morphology of staphyla and gongylidia

Using samples collected directly from the fungus gardens of live leafcutter colonies, we first used scanning electron microscopy (SEM) to visualize the external morphology of gongylidia and staphylae. The samples were fixed in PBS with 0.1% Tween 20 and fixatives (4% glutaraldehyde, 4% formaldehyde) and then dehydrated in an ethanol series (35%, 55%, 75%, 85%, 95%, and 2x in 100%) for 30 minutes per concentration, critical-point dried, coated with platinum, and imaged on a JSM-840 scanning electron microscope (JEOL, Tokyo, Japan) at 7.0 kV at the Zoological Museum of the University of Copenhagen. A slight wrinkled appearance of the surface of gongylidium cells in the resulting SEM images was due to unavoidable plasmolysis caused by the preparation process.

We then used light, fluorescence and confocal microscopy to view the cultivar’s internal morphology (*e.g.* septa, vacuoles, nuclei, etc.) and examine cellular reorganizations associated with gongylidium induction. For all imaging, staphylae were first placed in a drop of mounting solution (dH_2_O, PBS, 3% KOH) on a glass slide. Gongylidia were then separated under a stereo microscope (16x or 25x magnification) with 0.16-mm diameter acupuncture needles and stained. For visualization with white light, we placed samples in either 0.1% Congo-red (in 150 mM NaCl) for one minute followed by a wash with 150 mM NaCl, or 1.5% phloxine followed by a wash with 3% KOH. For nucleus visualization under UV light, we stained samples for 10 minutes using “Vectashield with DAPI” (Vector Laboratories, Burlingame, CA, USA). For confocal imaging, we stained the staphylae with dextran conjugated with Alexa Fluor 647 (Invitrogen, MA, USA) in the concentration of 100 μg/ml in PBS for 30 minutes and washed twice in PBS. We then acquired images at magnifications of up to 400x by performing bright-field, dark-field, phase-contrast, and fluorescence microscopy under an Olympus BX63 microscope (Olympus, Tokyo, Japan). The microstructures were measured and photographed using a mounted QImaging Retiga 6000 monochrome camera and cellSens Dimension v1.18 (Olympus) image-processing software. Confocal images were acquired with a Dragonfly spinning-disk confocal system (Andor Technology, Belfast, Northern Ireland) equipped with a 25x water-immersion objective. Samples were excited with a 637 nm laser line and fluorescence was collected with a 698/77 emission filter. Images were processed using FIJI software.

We next used transmission electron microscopy (TEM) to visualize fungal cells with greater magnification and resolution (i.e. the 500 nm scale). Staphylae were collected from fragments of intact fungus gardens, fixed in 2% glutaraldehyde in 0.05 M PBS (pH 7.2) and then post-fixed in 1% w/v OsO_4_ with 0.05M K_3_Fe(Cn)_6_ in 0.12 M sodium phosphate buffer (pH 7.2) for 2 hr at room temperature. Fixed samples were washed three times in ddH_2_O for 10 minutes and dehydrated in a series of increasing ethanol concentrations series (70%, 96% and 99.9%). Each dehydration lasted 15 min and was performed twice per concentration. Samples were then repeatedly infiltrated for 20 to 40 min with increasing Resin Epon:Propylene oxide ratios (1:3, 1:1, 3:1) and subsequently embedded in 100% Epon and polymerized overnight at 60°C. Sections of 60 nm thickness were then cut with an Ultracut 7 ultramicrotome (Leica, Vienna, Austria), collected on copper grids with Formvar supporting membranes, and stained with both uranyl acetate and lead citrate. These samples were TEM imaged on a CM100 BioTWIN (Philips, Eindhoven, The Netherlands) at an accelerating voltage of 80 kV. Digital images were recorded with a side-mounted OSIS Veleta digital slow scan 2 × 2 k CCD camera and the ITEM software package (Olympus Soft Imaging Corp, Münster, Germany). This sample preparation and imaging was performed at the Core Facility for Integrated Microscopy at the University of Copenhagen.

### Autophagy inhibition assay

We tested the role of autophagy in gongylidium production using an *in vitro* growth assay with four treatment groups. Autophagy is often initiated in cells when the target of rapamycin kinase (TOR) is inhibited. Autophagy can thus be induced *in vitro* by adding rapamycin (RAP) [50] an allosteric TOR inhibitor [48]. Autophagy is often inhibited *in vitro* using Chloroquine (CQ) or 3-methyladenine (3-MA), as these compounds respectively block autophagosome-vacuole fusion [51] and suppress an enzyme (class III PtdIns3K) required to initiate autophagosome formation [52]. We compared cultivars grown in the dark for 46 days at 25°C on a baseline Potato Dextrose Agar (PDA) medium containing nutrients known to maximize cultivar performance [19, 53] (n = 40) with cultivars grown on plates with RAP (n = 27), CQ (n = 28), or 3-MA (n = 30). Briefly, 5-mm diameter fungus plugs from previously isolated and reinoculated PDA culture were placed in 60-mm Petri dishes containing 10 ml of PDA (control), PDA + 300 ng/ml rapamycin (Medchem Express, Monmouth Junction, NJ, USA), PDA + 1.5 mM chloroquine diphosphate (Sigma-Aldrich, St. Louis, Missouri, USA), or PDA + 10 mM 3-MA (Medchem Express). We then photographed plates to measure growth area (mm^2^) using ImageJ [54] and directly counted the staphylae on these Petri dishes under a stereo microscope (40x magnification) to quantify staphyla density (number of staphylae/growth area). The measurement of growth area also enabled assessment of other unintended inhibitory effects of the added chemicals on cultivar performance. We tested for treatment effects (PDA-Control, RAP, CQ, 3-MA) on mycelial growth and staphyla density in R version 4.0.2 [55] using a Kruskal-Wallis test in *rstatix* version 0.7.0 [56] with pairwise post-hoc tests performed using Dunn’s Test in *rstatix* with adjusted p-values calculated using the false-discovery rate method.

### Transcriptome sequencing, assembly and differential expression analyses

To detect whether upregulated transcripts in staphylae were enriched with autophagy genes, we first collected staphylae and undifferentiated mycelia from axenic fungal cultures isolated from an *A. colombica* colony (Ac2012-1) and grown on PDA medium for 30 days as specified above. From individual Petri dishes, we then used a RNeasy plant mini kit (Qiagen, Germany) to extract total RNA from differentiated staphyla (pooled 200 staphylae individually collected with sterile acupuncture needles, n = 5 Petri dishes samples) and undifferentiated mycelia (adjacent mycelia lacking staphylae, n = 5 Petri dishes samples). Samples were immediately placed in Qiagen RLC buffer containing 10uM DTT and RNA was then extracted using the manufacturer-specified protocol. These samples were sent to BGI Europe (Copenhagen, Denmark) where mRNA enrichment with oligo dT and strand-specific libraries were constructed using dUTP in the cDNA synthesis. For each sample, between 24 and 30 million clean 100bp paired-end reads were generated by a DNBseq-G400 sequencer (MGI Tech, Shenzhen, China).

We used pooled clean reads from all samples (staphylae and mycelia) to assemble a *de novo* transcriptome using Trinity-v2.12.0 [57] and default settings with the addition of Jaccard-clip and strand-specific (SS) options, to reduce the generation of chimeric transcripts and account for the SS library construction, respectively. All downstream tools accounted for SS sequences (analyzing only positive strands or sequences). Using CD-HIT [58] to cluster highly similar sequences (98% similarity), we reduced the assembly from 93,470 to 78,820 transcripts ranging from 187 to 26,203 bp. We used Trinity built-in pipelines to first estimate transcript abundance with RSEM v1.3.1 [59] and then build a transcript expression matrix. We then performed a differential expression analysis with a trinity built-in pipeline using DESeq2 [60] setting the analysis to filter for the differentially expressed transcripts (DETs) with log_2_ fold change > 1.0 and *P* < 0.001. To annotate the DETs, we first converted transcripts to amino acid sequences using Transdecoder (https://github.com/TransDecoder/TransDecoder) to identify the longest open read frames (orfs) per transcript and translate them into amino acid sequences, and then annotated the DETs with KEGG database [61] using the online tools BlastKOALA, GhostKOALA [62] and KofamKOALA [63].

## Data availability

All the generated RNA-seq datasets are available at NCBI under the BioProject ID PRJNAXXXXXX.

## Acknowledgements

This study was funded by a European Research Council Starting Grant (ELEVATE: ERC-2017-STG-757810) to JZS. Benjamin Conlon assisted with statistical analyses. Useful comments were provided by Jacobus Boomsma, Gareth W. Griffith and Pedro Elias Marques. The images recreating stages of gongylidium formation in Figure 4 were produced by Damond Kyllo. Assistance with transmission and scanning electron microscopies was provided by the Core Facility for Integrated Microscopy, Faculty of Health and Medical Sciences, and the Zoological Museum, both at University of Copenhagen. Data processing and analysis were performed using Computerome, the National Life Science Supercomputer at DTU (www.computerome.dk). Sylvia Mathiasen and Rasmus Larsen provided general laboratory assistance.

## Supplementary figures

**Fig. S1:**
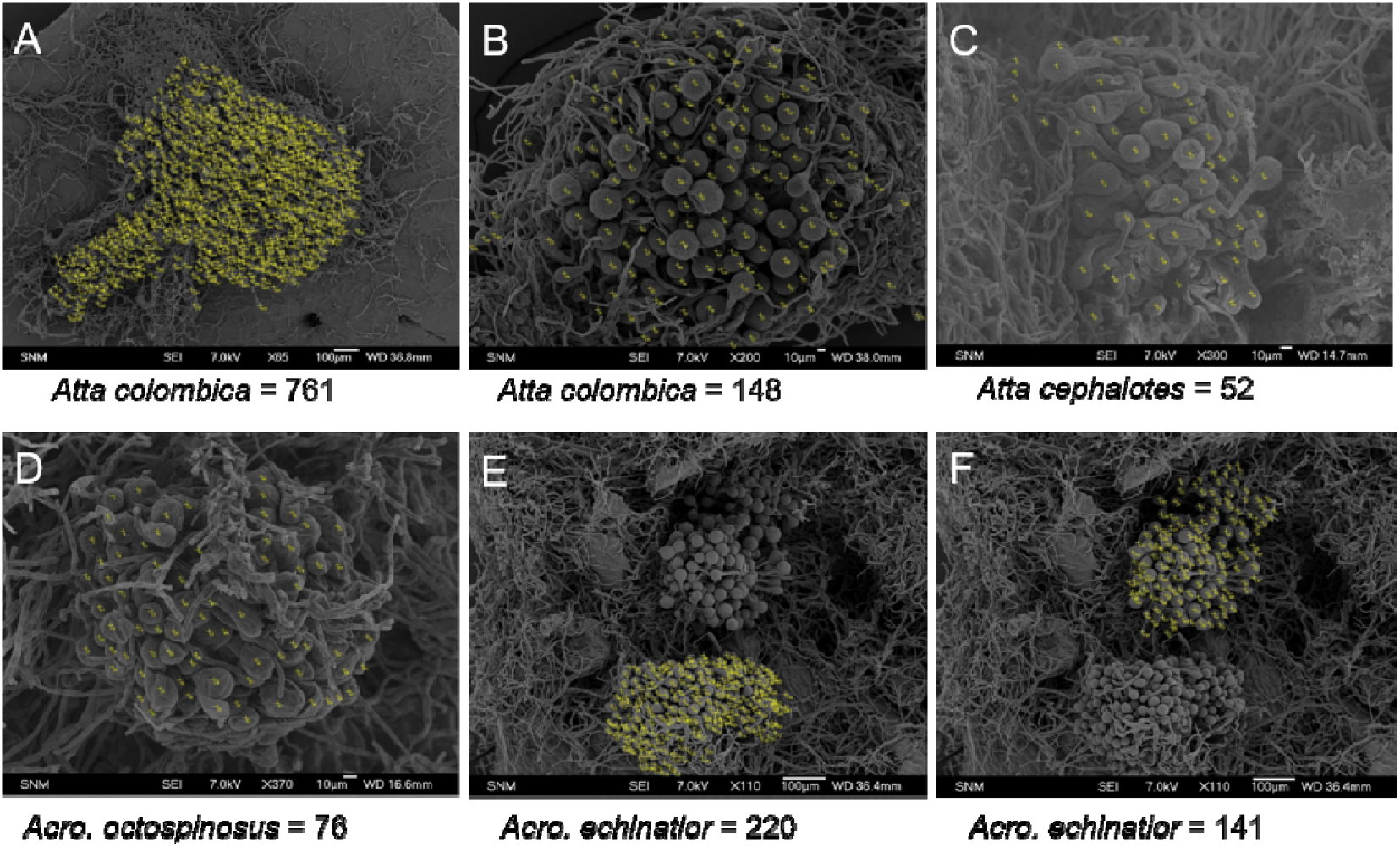
Representative counts of gongylidia per staphylae in *L. gongylophorus* from different leafcutter ants’ species. A) Staphylae from in vitro culture without ant manipulation. B-E) staphylae from colonies fungus garden. Yellow marks indicate individual gongylidia count on ImageJ. Scale bars sizes are indicated in each image.

**Fig. S2:**
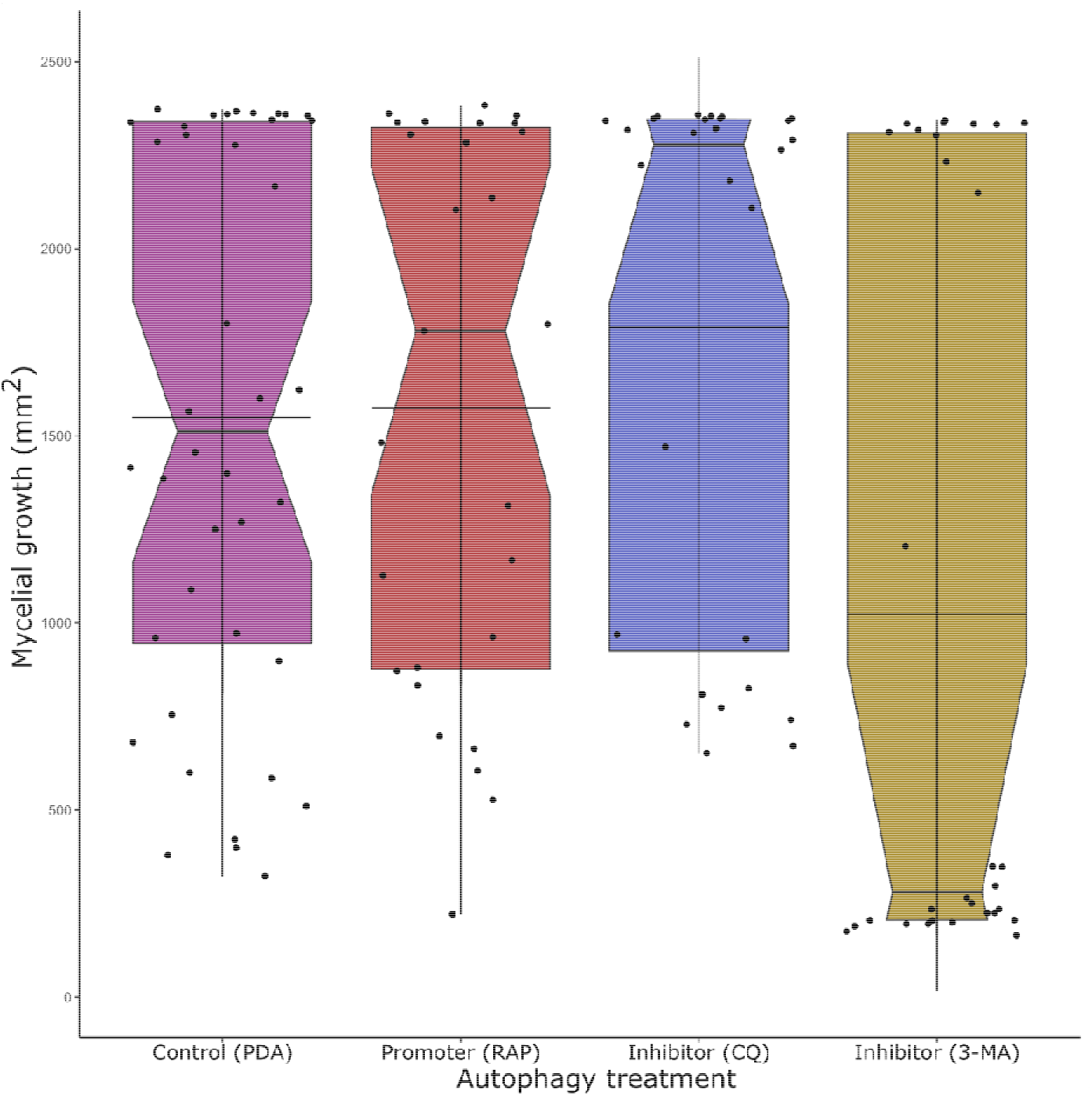
Mycelial growth distribution (area per treatment). The mycelial growth showed significant differences between 3-MA and all other treatments (PDA:3-MA, *p_adj_* = 0.004; RAP:3-MA, *p_adj_* = 0.015; CQ:3-MA, *p_adj_* = 0.001), but no other significant pairwise comparisons. Thus, while both autophagy inhibition treatments resulted in staphyla reduction, it is possible that 3-MA negatively influenced staphyla density through unknown indirect effects on cultivar performance. Horizontal bars indicate the distribution means.

**Fig. S3:**
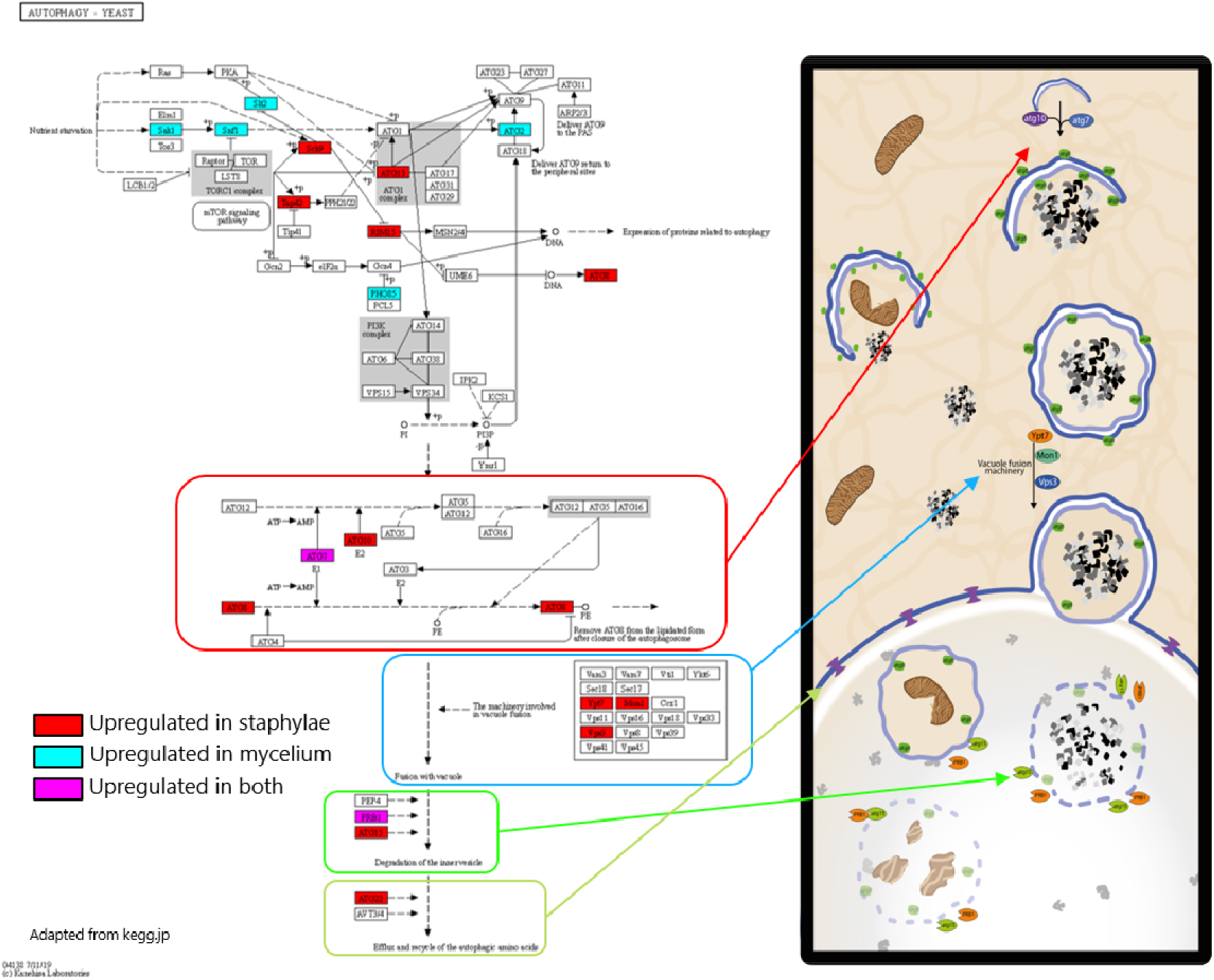
Autophagy metabolic pathway map displaying gene expression in *L. gongylophorus*. Highlighted genes show upregulated transcripts in staphylae (red), mycelium (cyan) or in both tissues (magenta). The steps of the pathway are illustrated on the right showing where the products of these genes will act during autophagy. Adapted from KEGG website (www.kegg.jp).

**Video S1: Time lapse of staphyla stained with dextran-Alexa Fluor 647 under confocal microscope over 20 minutes.** Red arrows point gongylidia in which is possible to observe autophagic bodies trapped within the vacuole. Some of these vesicles are filled with dextran and appear more fluorescent than the vacuole lumen, others are filled with non-fluorescent material and appear darker than the vacuole.

